# Iterative Cochran’s C test as a multivariate method to detect higher than expected variability: a microbiological inter-laboratory ring trial as a case-study

**DOI:** 10.1101/2020.11.19.389619

**Authors:** Antonio Monleon-Getino, AC Marca Home Care

## Abstract

**Introduction:** An interlaboratory calibration analysis was carried out to validate a methodology for a European standard for domestic laundry disinfection, using different doses of disinfectant and microorganisms. ISO 5725-2 and ISO 13528 form the basis of interlaboratory validations of quantitative methods, but there is a need for a simple graphical method to detect differences in laboratory behavior in terms of accuracy and variability.

**Objective:** A novel multivariate method based on the classical <u>Cochran’s C test</u>, as well as PCA and bootstrapping, which allows the inclusion of different correlated variables, was applied to identify higher variability than expected in factor (e.g. laboratories) levels, and the detection of multivariate outliers in a reduced space.

**Methods:** The proposed method is based on resampling, using the same sample many times but removing cases at random and performing Cochran’s C test for all the variables together in a reduced space.

**Results:** The method was tested by checking 7 laboratories for high variability in different parameters (logarithmic reduction (LR), cross contamination (RI), and wash water (WW)). After applying the proposed statistical analyses, no reasons were found to reject any of the participating laboratories. Multiple applications of the method are possible and we describe a case study in which the multivariant iterative <u>Cochran’s C test</u> was used: variability detection with multiple microbiological parameters (with high variability) during an interlaboratory ring trial.

## 1. Background

The high number of uncontrollable variables in microbiological systems increases experimental complexity and reduces accuracy, which can lead to the misinterpretation of data or uncorrectable errors [1][2][3].

The great number of variables at play in each microbiological study arise from the inherent complexity of living systems. Each variable contributes a certain degree of error which, depending on the system, may propagate linearly or non-linearly. The more variables involved, the more errors can be expected, which can modify the outcome of a given study. The causes of variability in microbiology methods can be roughly divided into three groups: the test system (e.g. microorganism and environmental conditions), the scientist(s) performing the study, and in the case of antimicrobial efficacy studies, the test substance (1). Although a number of sources of variability have been defined, there may be others of equal significance unknown to microbiologists (1). In general, experimental results must be reproducible to be meaningful. As variability is known to exist in biological systems and is impossible to avoid, strategies should be sought to reduce its impact on significance.

In the field of textile disinfection [3], there are standard laboratory practices (as EN16616) that use microbiological systems to validate the efficacy of a disinfectant product under simulating use conditions (P2S2 tests). These tests, which are designed for hospital areas, use industrial washing machines. A new recently developed technique for testing domestic disinfectants uses a laboratory-scale device to simulate the mechanical effect of the wash. A ring trial was performed to test the validity of the new approach in terms of accuracy and variability and the data was analysed according to ISO 5725-2 and ISO13528 and the new statistical method.

The accuracy and variability of a reference test are critical to establish the performance of a new product. During the ring trial, reductions of a known microbial population exposed to the product during a wash cycle and a rinsing cycle was evaluated. Microbiological variables like LR (logarithm reduction log (CFU/ml) for different types of microorganisms), WW (counts of microorganisms in the wash water) or RI (recovery of microorganisms in the cross-contamination carrier) were generated and compared statistically between laboratories, laboratory-scale devices, product and product dose.

The repeatability and reproducibility of the method when tested with water was calculated from the collected data.

### Interlaboratory methods and detection of variability

The library ILS [4] for R can be applied in interlaboratory studies to detect laboratories that provide inconsistent results compared to the others. It allows various testing materials to be used simultaneously, using standard univariate and functional data analysis (FDA) perspectives. The univariate approach based on ASTM E691-08 [3][4] consists of estimating Mandel’s h and k statistics to identify those laboratories producing the most significantly different results; the presence of outliers is determined by Cochran’s and Grubbs tests, and analysis of variance (ANOVA) techniques are provided (F and Tuckey tests) to test for differences in means corresponding to different laboratories per material. The functional nature of data retrieved in analytical chemistry, applied physics and engineering (spectra, thermograms, etc.) is taken into account. The ILS package also provides an FDA approach to determine Mandel’s k and h statistics distribution by smoothed bootstrap resampling [4].

For another example of intercalibration analysis for biological data, see: http://www.nmbaqcs.org/media/1676/phy-icn-16-mi1-final-report-vr1.pdf

**Table 1:**
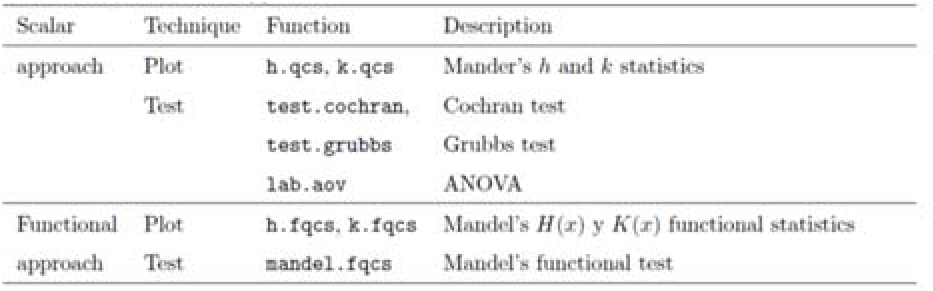
Some of the tests used to check the current interlaboratory consistency analysis recommended by Flores et al. (2018) [4]

Most of this hypothesis test is included in ISO 5725–2:1994 (E) [5] [6] Accuracy of measurement methods and results – Part 2: Basic method for the determination of repeatability and reproducibility of a standard measurement method (See https://www.aenor.com/normas-y-libros/buscador-de-normas/iso/?c=011834)

## 2. Objectives

We aimed to apply a new multivariate method to detect higher than expected variability in factor levels in cases with multiple attributes, using PCA and boostrap. The method was tested in an interlaboratory ring trial to detect laboratories with higher-than-expected sets of microbiological variables.

## 3. Material and methods

This new method is based on a previous study by Monleón & Ocaña (2019) [7].

The method proposed is based on resampling (performing 10,000 iterations, where samples are removed and bootstrapped), using the same sample many times but removing cases at random and performing Cochran’s C test for all the variables together in a reduced space. After carrying out the statistical tests, it is possible to count how many times they were significant. If frequent, it suggests that the laboratory may have an issue with the variability of its samples.

### Classical Cochran’s C test

Cochran’s C test is used to test the assumed homogeneity of variance among the residuals in the analysis of variance. The test statistic is a ratio that relates the largest empirical variance of a particular treatment with the sum of the variances of the other treatments.

In the present study, Cochran’s C test was applied to decide if a single estimate of laboratory variance is significantly larger than a group of laboratory variances with which the single estimate is supposed to be comparable. In this exercise, various parameters were evaluated simultaneously (LR, RI and WW). Furthermore, Cochran’s C test assumes a balanced design (equal size of samples per laboratory) and assumes that each individual data series is normally distributed, which is not a common situation [8].

Cochram’s C test detects one exceptionally large laboratory variance value at a time. The corresponding data series is then omitted from the full data set. According to ISO standard 5725 [<u>ISO</u> Standard 5725-2:1994] [5], Cochram’s C test may be iterated until no further exceptionally large variance values are detected, but this may lead to excessive rejections if the underlying data series are not normally distributed.

The C test tests the null hypothesis (H_0_) against the alternative hypothesis (H_1_):

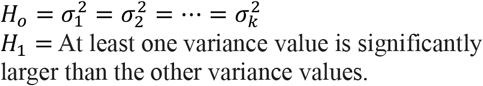

For *k* laboratories

the Cochram test is based on the the ratio *C_j_*:

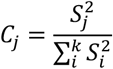

where:

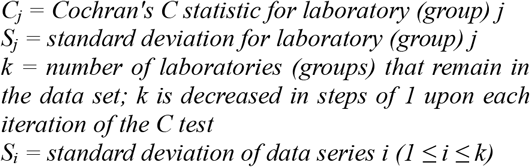

<u>Cochran’s C</u> test was transformed to be interactive by random resampling of the samples and using all the v variables at the same time. In order to observe extremely high inter-batch variabilities using <u>Cochran’s C</u> test for all the v variables (e.g. LR, RI, WW) considered together, we propose a multivariant version with two strategies: 1) the test is performed for each of the v variables considered, performing 10000 (or more) resamplings (with substitution) of the data set (only reference batches) and scoring (%) how many times the <u>Cochran’s C test</u> is significant (p <0.05) for each of the k variables, and 2) a PCA is carried out with the set of k variables, taking the first and second component (PC1, PC2) and then repeating step 1). The result is evaluated based on the score obtained, so if the percentage of times is greater than a certain value (we suggest >40%) for the total of k variables, the elimination of that batch will be considered for the bioequivalence analysis.

The procedure was encapsulated in the function:

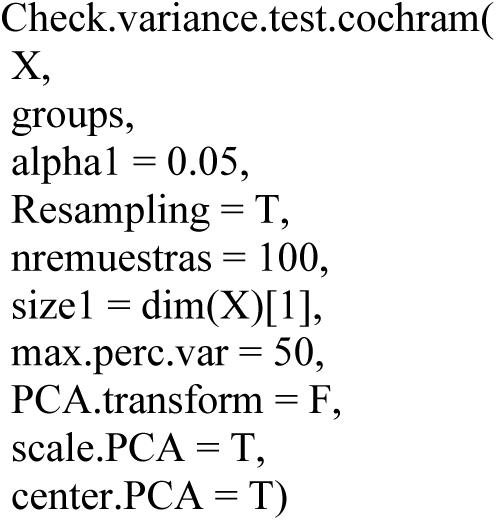

Check.variance.test.cochram() function is part of the R library called Diagnobatch () developed by Monleón (2020) [9]. Although some of these methods are novel in microbiology, they have been widely used in other industrial sectors, such as the pharmaceutical industry [7]. In situations like the one studied here, a large number of redundant variables are involved and accuracy, variability and outlier hypotheses must be verified (using the common statistical methods: Cochran’s C test, Fligner-Killeen test, Grubbs test) in order to detect inconsistencies between groups (in this case laboratories), but using a multivariate approach and summarizing the variables through principal components analysis [7].

## 4. Results

The proposed method was applied during an interlaboratory ring trial to check if there was high variability between 7 laboratories simultaneously evaluating several microbiological variables.

In the first of two situations (Figure 1), after gathering all the LR type variables for the different microorganisms using PCA and performing the modified <u>Cochran’s C test</u>, nothing remarkable was detected in terms of higher than normal variability between laboratories.

**Figure 1:**
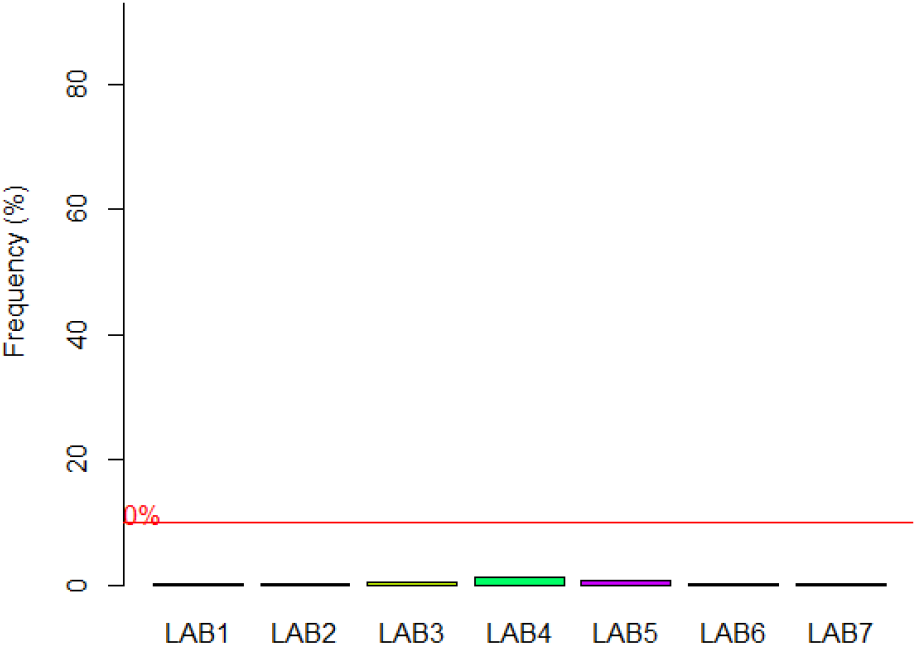
Plot bar showing the variability between laboratories for all LR variables included. The bars represent the percentage of times the test was significant (p<0.05) in 10000 applications of Cochran’s C test.

In a second situation (Figure 2), another group of variables related to **RITSA, RIMEA, WWTSA, WWMEA** was used and the dimension was reduced by means of PCA. In this case, variances higher than normal were detected for one of the levels tested.

**Figure 2:**
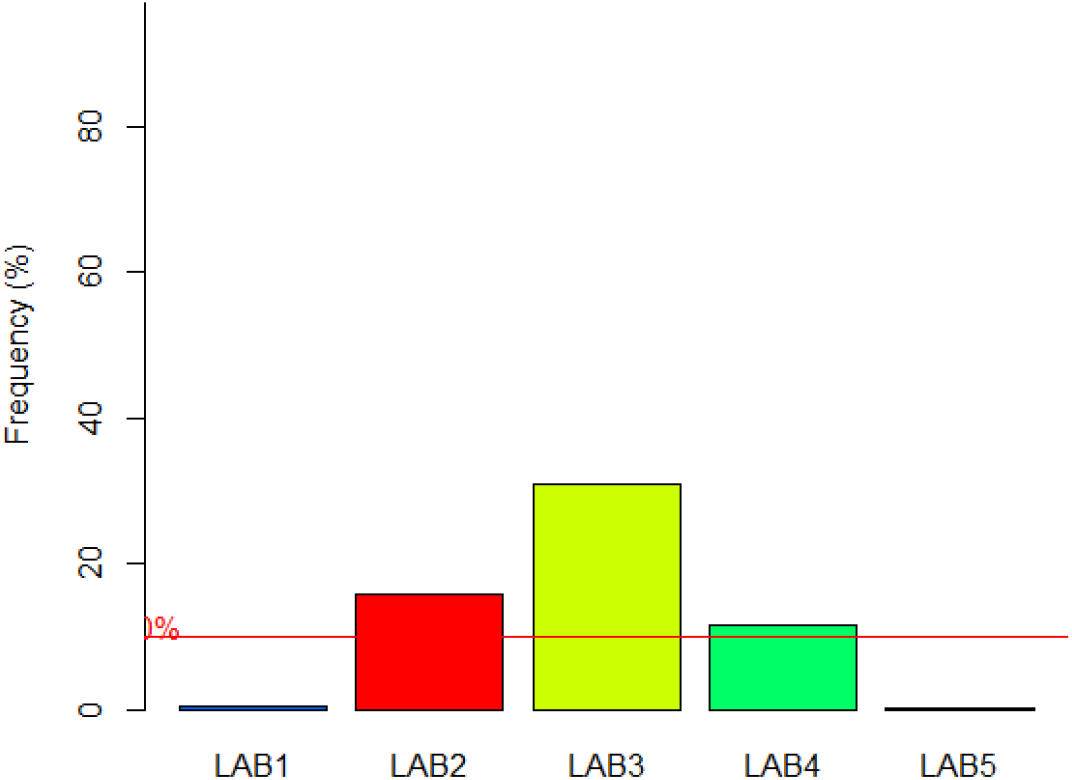
Plot bar showing the variability between laboratories when all variables (RITSA, RIMEA, WWTSA, WWMEA) are included. The bars represent the percentage of times the result was significant (p<0.05) in 10000 applications of Cochran’s C test. Yellow and red laboratories are notable.

As shown in Figure 1, significant differences in variances were detected in only a few tests (1-2%), and laboratories with an abnormal variability can be discarded.

As depicted in Figure 2, laboratory 3 stands out in presenting higher than normal variances (more than 35% of 10,000 tests) according to the <u>Cochran’s C test</u> for multiple variables. Nevertheless, there is no value (cut-off or critical or p-value) from which it can be said that this is a problem because the number of times of the Cochran C test is significant is bellow 50%. If the number was above 50%, we could affirm that the variable was higher than expected, and investigate the reasons why. Therefore, in this example, no reason was found to reject any laboratory.

## 5. Conclusions

This study presents a novel version of the classical Cochran’s C test that can detect higher than expected variability in factor levels in cases with multiple attributes using PCA and boostrapping. By using random resampling of samples and all the v variables at the same time, <u>Cochran’s C test</u> was transformed to be interactive.

During an interlaboratory calibration analysis to validate a methodology that will be proposed as a European standard for domestic laundry disinfection, some variables had a higher than expected variability among the laboratories. This problem served as an opportunity to test the developed method.

It is difficult to decide if a laboratory has a problem of excessive variability when multiple microbiological variables of great intrinsic variability are being used.

Different results are presented with real microbial data to validate the proposed method. In one case, a higher than expected variability was ruled out and in a second case, a laboratory with a higher than expected variability was detected.

Multiple applications of the proposed method are possible in other fields of industry and science.

## Notes

### Competing Interest Statement

The authors have declared no competing interest.

